# Quantification of permethrin resistance and *kdr* alleles in Florida strains of *Aedes aegypti* (L.) and *Aedes albopictus* (Skuse)

**DOI:** 10.1101/365171

**Authors:** Alden S. Estep, Neil D. Sanscrainte, Christy M. Waits, Sarah J. Bernard, Aaron M. Lloyd, Keira J. Lucas, Eva A. Buckner, Rajeev Vaidyanathan, Rachel Morreale, Lisa A. Conti, James J. Becnel

**Affiliations:** Navy Entomology Center of Excellence, CMAVE Detachment, Gainesville FL; United States Department of Agriculture, Agricultural Research Service, Center for Medical, Agricultural, and Veterinary Entomology, Mosquito and Fly Research Unit, Gainesville FL; Pasco County Mosquito Control District, Odessa, FL; Collier County Mosquito Control District, Naples, FL; Manatee Mosquito Control District, Palmetto FL; Clarke Inc., Saint Charles, IL; Lee County Mosquito Control, Lehigh Acres, FL; Florida Department of Agriculture and Consumer Services, Tallahassee, FL

## Abstract

Recent outbreaks of locally transmitted dengue and Zika viruses in Florida have placed more emphasis on the importance of integrated vector management plans for *Aedes aegypti* (L.) and *Aedes albopictus* Skuse. Adulticiding, primarily with pyrethroids, can be the best option available for the immediate control of potentially arbovirus-infected mosquitoes during outbreak situations. While pyrethroid resistance is common in *Ae. aegypti* worldwide and testing is recommended by CDC and WHO, resistance to this class of products has not been widely examined or quantified in Florida. To address this information gap, we performed the first study to quantify both pyrethroid resistance and genetic markers of pyrethroid resistance in *Ae. aegypti* and *Ae. albopictus* strains in Florida. Using direct topical application, we examined 21 *Ae. aegypti* strains from 9 counties and found permethrin resistance (resistance ratio (RR)=6-61-fold) in all strains when compared to the susceptible ORL1952 control strain. Permethrin resistance in five strains of *Ae. albopictus* was very low (RR<1.6) even when collected from the same containers producing resistant *Ae. aegypti*. Characterization of two sodium channel *kdr* alleles associated with pyrethroid-resistance showed widespread distribution in 62 strains of *Ae. aegypti*. The 1534 phenylalanine to cysteine (F1534C) single nucleotide polymorphism SNP was fixed or nearly fixed in all strains regardless of RR. We observed much more variation in the 1016 valine to isoleucine (V1016I) allele and observed that increasing frequency of the homozygous V1016I allele correlates strongly with increased RR (Pearson corr= 0.905). In agreement with previous studies, we observed a very low frequency of three *kdr* genotypes, IIFF, VIFF, and IIFC. In this study, we provide a statewide examination of pyrethroid resistance, and demonstrate that permethrin resistance and the genetic markers for resistance are widely present in FL *Ae. aegypti*. Resistance testing should be included in an effective management program.

**Author Summary:** *Aedes aegypti* and *Aedes albopictus* can vector a variety of arboviruses that cause diseases and are thus a public health concern. Pyrethroid insecticide resistance is common in *Aedes aegypti* in many locations worldwide and can adversely affect vector control operations. However, the resistance status of these vectors in Florida is largely unreported and recent local transmission of dengue and Zika viruses has made this information critical for effective control operations. In this study, we showed that permethrin resistance and two common SNPs of the voltage gated sodium channel (V1016I and F1534C) previously associated with pyrethroid resistance were widely present in Florida *Aedes aegypti* strains. We also observed a strong correlation between the IICC genotype and RR as determined by topical application, which suggests, as have others, that *kdr* frequency may be a useful indicator of resistance in *Aedes aegypti*.

## Introduction

Local vector control programs play a part in public health. In many countries, these programs serve as the primary defense against the spread of several mosquito-borne diseases. Effective Integrated Vector Management (IVM) programs rely on surveillance information coupled with multiple vector control strategies, such as chemical adulticiding, when needed to reduce vector populations as well as arbovirus transmission. Limited recent transmission of locally acquired dengue and Zika viruses in the southeastern continental US, primarily Florida, has brought renewed attention to the importance of IVM programs where the potential vectors, *Aedes aegypti* (L.) and *Ae. albopictus* Skuse, have long been present. However, for IVM programs in Florida to effectively control *Aedes* vectors and to reduce dengue and Zika virus transmission during an outbreak, it is essential to know which adulticide products are effective against local *Ae. aegypti* and *Ae. albopictus* strains.

Organophosphates and pyrethroids are the only two classes of insecticides available for public health use to control adult mosquitoes in the US. Compared to organophosphate insecticides, pyrethroids have higher public acceptance, rapid knockdown, relatively low costs, and are generally the product class of choice when adulticiding is required [1] Unfortunately, years of pyrethroid insecticide use and previous DDT usage has increased the frequency of genetic and enzymatic resistance in insects like mosquitoes. Genetic target site changes of the sodium channel, known as knockdown resistance *(kdr)* mutations, are a relatively common insect response to selective pressure by pyrethroids [2,3]. Although a variety of other single nucleotide polymorphisms (SNPs) have been noted, in *Ae. aegypti*, the primary two SNPs linked to pyrethroid resistance are at codons 1016 and 1534 (positions according to standard *M. domestica* notation) [4,5]. One allele of the 1016 mutation is geographically distinct to the Western hemisphere and results in the replacement of the normal valine with an isoleucine (V1016I), while the 1534 mutation results in a phenylalanine to cysteine change (F1534C). There is currently debate about the individual toxicological effect of these two mutations, but they are consistently present in resistant strains [4]. Heterozygous and homozygous combinations of these two SNPs could result in nine possible genotypes, but strong linkage between the SNPs has been noted with only six of the genotypes observed in a study of Mexican *Ae. aegypti* [6].

Pyrethroid resistance and *kdr* alleles have been well documented in *Ae. aegypti* strains from Central America, South America and the Caribbean [7-11]. Recent testing as part of the Zika emergency response in Puerto Rico has shown that isolated, early reports of resistance were indicative of widespread resistance on the island [11-14]. Studies have also shown the pyrethroid resistance is not specific to a particular chemical but is generally class-wide to type I, type II, and non-ester pyrethroids [11, 15, 16]. *Kdr* alleles and pyrethroid resistance are widely distributed in Mexico, including Nuevo Laredo which lies just south of the US-Mexico border [9].

In contrast, little has been published about pyrethroid resistance in *Aedes* strains within the continental US. In the Garcia et al. [9] study mentioned above, the authors did not find *kdr* alleles in a Houston, TX collection during 1999. Two recent reviews of the *Aedes* resistance literature listed no reports of resistance in continental US *Ae. aegypti* [4, 17], but Cornel et al. [18] recently demonstrated toxicological resistance and sodium channel mutations in invasive populations of *Ae. aegypti* in California. Two recent studies using CDC bottle bioassays do indicate resistance in strains from the southern US, but these studies do not provide any quantification of the strength of the resistance nor did they examine the presence of *kdr* alleles [19, 20]. In contrast, *Ae. albopictus* does not appear to frequently develop *kdr* resistance, and most US strains tested thus far have only shown very minimal pyrethroid resistance [4, 21, 22].

Resistance surveillance is recommended by the CDC and statewide initiatives to map pyrethroid resistance have begun in Florida and California [20,23]. Resistance information is critically important as strong pyrethroid resistance can cause failure of adulticiding control interventions. In this study, a collaborative group of government, academic, industry and vector control district stakeholders collected *Ae. aegypti* and *Ae. albopictus* adults, eggs or larvae from more than 200 locations throughout Florida to assess the extent and intensity of pyrethroid resistance. The goal of this study was to improve vector control operations by producing a resistance map and apply this information to make more effective control decisions. We used direct topical application of permethrin to determine resistance ratios (RRs) relative to the susceptible ORL1952 strain for 21 wild type strains of Ae. *aegypti* and 5 strains of *Ae. albopictus*. Nearly 5,000 *Ae. aegypti* from numerous locations were genotyped by allele-specific PCR to assess the frequency of V1016I and F1534C alleles and rapidly visualize the pattern of resistance throughout the state of Florida.

## Methods

### Mosquito strains and rearing

The toxicological profiles and rearing procedures for the susceptible (ORL1952) and resistant (Puerto Rico -BEIResources, NR-48830) strains of *Ae. aegypti* used in this study have been described previously [15]. Strain information, including specific location information, collectors, and dates of collection are noted in Table 1. Field strains for toxicology testing were often collected as mixed *(Ae. albopictus* and *Ae. aegypti)* eggs laid on seed germination paper (Anchor Paper, St. Paul, MN) placed in a variety of oviposition containers including black plastic stadium cups, plastic cemetery vases, and glass containers. Field collected eggs were hatched by soaking papers for 48-72 hours at 27°C in rearing trays of untreated well water or deionized water (diH_2_O). Papers were carefully removed and larvae were reared through adult emergence with a quarter ration of 3:2 larval food compared to the standard rearing method for the ORL1952 susceptible strain [24] due to sensitivity to overfeeding. Strains from Monroe, Seminole, Orange, Hernando and Sarasota counties were collected as larvae from a variety of manmade and natural containers (tires, plant pots, bottles, buckets, etc.), rinsed with diH2O and then placed into the standard larval rearing procedure described above. Pupae were collected from rearing trays and placed in emergence chambers or 12”x12” screen cages (BioQuip Models 1425 & 1450B, Rancho Dominquez, CA). After emergence, wildtype mosquitoes were briefly chilled to 4°C and sorted to species. Strains for toxicology testing were produced from locations that had more than 30 wildtype founders. If multiple nearby locations were combined to produce a strain, the GPS location in Table 1 represents the GPS centroid of the sites that contributed the mosquitoes. Individual oviposition cup locations and life stages tested are provided in S1 File. Eggs were produced using standard rearing methods [15]. Colony strains were provided 2-10 blood-meals on weekly intervals to collect F1 eggs using warmed bovine blood. If feeding was poor, warmed blood was spiked with 1mM ATP as a phagostimulant. Bovine blood for mosquito feeding was purchased by the USDA under a contract with a local, licensed abattoir. Blood was collected during normal operations of the abattoir from the waste stream after animal slaughter. Under CFR9, Parts 1-3, tissues, including blood, collected from dead livestock intended as food are exempt from IACUC regulation.

**Table 1.** Collection information for Florida *Aedes aegypti* and *Ae. albopictus* used in this study.

| County | Strain | Species | GPS Centroid | Collection Date | Collector |
| --- | --- | --- | --- | --- | --- |
| Brevard | South Brevard | *Ae. aegypti* | -80.672, 28.055 | 6/9/2016 | J. S. |
| Broward | Deerfield-Tivoli | *Ae. aegypti* | -80.115, 26.307 | 6/7/2017 | L. S. |
| Broward | Deerfield-Cocoa Cay | *Ae. aegypti* | -80.099, 26.307 | 6/7/2017 | L. S. |
| Broward | Fort Lauderdale | *Ae. aegypti* | -80.181, 26.169 | 6/7/2017 | L. S. |
| Collier | Naples Park | *Ae. aegypti* | -81.809, 26.262 | 6/8/2017 | R. Ba |
| Collier | Old Naples | *Ae. aegypti* | -81.798, 26.141 | 6/8/2017 | R. Ba |
| Collier | Golden Gate City | *Ae. aegypti* | -81.695, 26.188 | 6/8/2017 | R. Ba |
| Collier | East Naples | *Ae. aegypti* | -81.768, 26.125 | 6/8/2017 | R. Ba |
| Duval | Aberdeen | *Ae. aegypti* | -81.701, 30.303 | 9/1/2016 | C. W. |
| Duval | Jean St. | *Ae. aegypti* | -81.710, 30.298 | 9/1/2016 | C. W. |
| Hernando | Brooksville | *Ae. aegypti* | -82.553, 28.450 | 7/1/2016 | Sanscrainte |
| Hernando | Brooksville | *Ae. albopictus* | -82.390, 28.557 | 7/1/2016 | Sanscrainte |
| Hillsborough | Marti Cemetery | *Ae. aegypti* | -82.494, 27.966 | 7/1/2014 | Estep/Be |
| Indian River | Vero Beach | *Ae. aegypti* | -80.419, 27.641 | 3/1/2016 | R. S. |
| Lee | Alva | *Ae. aegypti* | -81.609, 26.715 | 4/5/2016 | R. Mor |
| Lee | Alva | *Ae. albopictus* | -81.609, 26.715 | 4/5/2016 | R. Mor |
| Lee | Cen. Cape Coral | *Ae. aegypti* | -81.945, 26.562 | 4/5/2016 | R. Mor |
| Lee | Cen. Lehigh Acres | *Ae. aegypti* | -81.626, 26.610 | 4/5/2016 | R. Mor |
| Lee | Fort Myers | *Ae. aegypti* | -81.887, 26.617 | 4/5/2016 | R. Mor |
| Lee | Banyan Creek | *Ae. aegypti* | -81.966, 26.502 | 4/5/2016 | R. Mor |
| Lee | San Carlos | *Ae. aegypti* | -81.820, 26.472 | 4/5/2016 | R. Mor |
| Manatee | Palmetto | *Ae. aegypti* | -82.559, 27.550 | 4/6/2016 | E. |
| Manatee | Cortez | *Ae. aegypti* | -82.685, 27.468 | 4/6/2016 | E. |
| Manatee | South Bradenton | *Ae. aegypti* | -82.663, 27.459 | 4/6/2016 | E. |
| Manatee | Longboat Key | *Ae. aegypti* | -82.682, 27.437 | 4/6/2016 | E. |
| Manatee | Anna Maria Island | *Ae. aegypti* | -82.739, 27.533 | 4/6/2016 | E. |
| Martin | Jensen Beach | *Ae. aegypti* | -80.227, 27.245 | 8/9/2017 |  |
| Miami-Dade | North Little River | *Ae. aegypti* | -80.207, 25.843 | 12/2016-01/2017 | C. |
| Miami-Dade | Central Little River | *Ae. aegypti* | -80.202, 25.840 | 12/2016-01/2017 | C. |
| Miami-Dade | Central Little River | *Ae. albopictus* | -80.202, 25.840 | 12/2016-01/2017 | C. |
| Miami-Dade | South Little River | *Ae. aegypti* | -80.206, 25.836 | 12/2016-01/2017 | C. |
| Miami-Dade | Miami Beach | *Ae. aegypti* | -80.136, 25.807 | 10/12/2016 | C. |
| Miami-Dade | North Wynwood | *Ae. aegypti* | -80.198, 25.808 | 10/11/2016 | C. |
| Miami-Dade | Central Wynwood | *Ae. aegypti* | -80.200, 25.801 | 10/11/2016 | C. |
| Miami-Dade | Central Wynwood | *Ae. aegypti* | -80.200, 25.801 | 10/11/2016 | C |
| Miami-Dade | East Wynwood | *Ae. aegypti* | -80.191, 25.801 | 10/11/2016 | C |
| Miami-Dade | Homestead I | *Ae. aegypti* | -80.475, 25.475 | 12/1/2016 | C |
| Miami-Dade | Homestead II | *Ae. aegypti* | -80.452, 25.459 | 12/1/2016 | C |
| Miami-Dade | Coconut Grove | *Ae. aegypti* | -80.233, 25.736 | 1/2/2017 | C |
| Miami-Dade | Coral Gables | *Ae. aegypti* | -80.269, 25.719 | 11/12/2016 | C |
| Miami-Dade | Aventura | *Ae. aegypti* | -80.140, 25.958 | 2/1/2017 | C |
| Miami-Dade | Hialeah | *Ae. aegypti* | -80.291, 25.853 | 1/2/2017 | C |
| Miami-Dade | Miami Lakes | *Ae. aegypti* | -80.314, 25.908 | 1/2/2017 | C |
| Miami-Dade | Kendall | *Ae. aegypti* | -80.353, 25.681 | 3/5/2017 | C |
| Miami-Dade | Doral | *Ae. aegypti* | -80.367, 25.811 | 1/2/2017 | C |
| Miami-Dade | Sunny Isles | *Ae. aegypti* | -80.123, 25.945 | 1/3/2017 | C |
| Miami-Dade | Brickell | *Ae. aegypti* | -80.195, 25.762 | 4/1/2017 | C |
| Monroe | Big Coppitt Key | *Ae. aegypti* | -81.663, 24.600 | 2/1/2016 | A. Estep/ D. J |
| Orange | Orlando | *Ae. aegypti* | -81.442, 28.551 | 5/1/2016 | K. Deutsc |
| Orange | Orlando | *Ae. albopictus* | -81.442, 28.551 | 5/1/2016 | K. Deutsc |
| Pasco | Port Richey-Stell | *Ae. aegypti* | -82.689, 28.257 | 6/8/2017 | A. Janusa |
| Pasco | Port Richey-Congress | *Ae. aegypti* | -82.707, 28.257 | 6/8/2017 | A. Janusa |
| Pasco | Port Richey-Pineland | *Ae. aegypti* | -82.706, 28.323 | 6/8/2017 | A. Janusa |
| Pasco | Port Richey-Gulf | *Ae. aegypti* | -82.690, 28.327 | 6/8/2017 | A. Janusa |
| Pasco | Holiday-Buena Vista | *Ae. aegypti* | -82.744, 28.184 | 6/8/2017 | A. Janusa |
| Pasco | Holiday-Scandia | *Ae. aegypti* | -82.738, 28.186 | 6/8/2017 | A. Janusa |
| Pasco | Elfers-Grove Park | *Ae. aegypti* | -82.733, 28.220 | 6/8/2017 | A. Janusa |
| Pasco | Elfers-Virginia City | *Ae. aegypti* | -82.710, 28.220 | 6/8/2017 | A. Janusa |
| Pasco | Zephyr Hills | *Ae. aegypti* | -82.182, 28.239 | 6/1/2014 | A. Llo |
| Pinellas | Clearwater | *Ae. aegypti* | -82.784, 27.948 | 9/1/2016 | J. |
| Sarasota | Sarasota | *Ae. aegypti* | -82.469, 27.213 | 9/1/2017 | W. Bren |
| Seminole | Altamonte Springs | *Ae. aegypti* | -81.406, 26.686 | 5/1/2016 | T. Jones |
| Seminole | Winter Park | *Ae. aegypti* | -81.323, 28.622 | 5/1/2016 | T. Jones |
| St. Johns | Lighthouse | *Ae. aegypti* | -81.314, 29.897 | 6/7/2016 | J. |
| St. Johns | Spanish Street | *Ae. aegypti* | -81.314, 29.896 | 6/7/2016 | J. |

GPS locations of strains represent the GPS centroid of multiple oviposition collections. Individual locations are listed in supplemental information.

*Aedes aegypti* strains from St. Augustine, Clearwater, and Vero Beach, FL were provided as F1 eggs produced at Anastasia Mosquito Control District, Pinellas County Mosquito Control, and the Florida Medical Entomology Laboratory, respectively.

To produce mosquitoes for bioassay testing, eggs from the F1 or F2 generation of field collected strains, the lab susceptible strains (ORL1952) and the pyrethroid-resistant strain were hatched and reared as described above in covered trays at a density of approximately 1000 mosquito larvae/tray. Adult mosquitoes were allowed to emerge into 12”x12” screen cages and provided with cotton saturated with 10% sucrose in diH_2_O. Females used for bioassays were 3-7 days post-emergence.

### Determination of resistance ratio by topical application

The adult topical bioassay has been described previously in detail [24, 25]. For these studies, the permethrin stock solution in DMSO (Product #N-12848-250MG, Chemservice, Westchester, PA) and all dilutions in acetone were prepared gravimetrically. An initial 10-fold dilution series was prepared over a range of relevant concentrations [15]. Sub-dilutions were prepared from the 10-fold dilution series as necessary to determine the critical region of the dose curve. Three assays (n=3) were performed for all strains with listed LD_50_s unless limited numbers of F1 test mosquitoes allowed only two replicates. Average mass/female was calculated for each strain by weighing a cohort of 50-100 females before each replicate.

LD_50_s, standard errors, and goodness of fit were determined from dose-mortality curves using SigmaPlot v13 (Systat software Inc., San Jose, CA) with data fit to a four parameter logistic model. To provide a comparative metric between strains that may have different body sizes, doses applied were divided by the average mass of the mosquitoes before curve fitting. This results in an LD_50_ of ng of active ingredient per mg of mosquito. Resistance ratios for strains were calculated using the LD_50_ of the field strain divided by the LD_50_ of the susceptible ORL1952 strain included in the same assay. In this study, we use the WHO scale to define the levels of resistance [26]. When RR is less than 5, the field population is considered susceptible. When the RR is between 5 and10, the mosquitoes are considered to have moderate resistance. A RR greater than 10 indicates that mosquitoes are highly resistant [27].

### Strain *kdr* genotyping

Genotypes for individual mosquitoes or eggs were determined using melt curve analysis with previously described allele-specific primers for the 1016 and 1534 SNPs [28, 29]. Assays were performed in 96 well plates on a StepOnePlus (Applied Biosystems) or QuantStudio5 (Applied Biosystems, Thermo Fisher Scientific, Waltham, MA) using SYBR green chemistry. Plates were loaded with 8μl of PCR master mix containing SYBR Select Master Mix (Applied Biosystems, Thermo Fisher Scientific, Waltham, MA), nuclease free water (NFW), and three balanced primers (Table 2). Primer balancing was necessary to ensure that the resulting melt curves would be accurately called by the analysis software. Individual adult females were homogenized for 60 seconds at max speed in 100μl of NFW on a bead beater (BioSpec, Bartlesville, OK). Individual eggs were similarly treated but homogenized in 30μl of NFW. Immediately after homogenization, samples were centrifuged at 10,000 relative centrifugal force (rcf) for 60 seconds to pellet solids. Two microliters of each supernatant were added to 8μl of PCR mix for each primer set and then subjected to standard cycling conditions (3min @ 95°C; 40 cycles @ 95°C for 3 sec, 60°C for 15 sec). Melt curve analysis followed cycling with acquisition of fluorescence data every 0.3°C as the temperature was ramped from 60°C to 95°C. Characteristic melting temperature (T_m_) peaks in the derivative fluorescence data indicate the presence of specific alleles [28, 29]. For the 1016 mutation, a codon for the susceptible valine has a T_m_ peak at 86±0.3°C while the isoleucine codon has a T_m_ of 77.3±0.3°C. The 1534 phenylalanine has a T_m_ of 79.8±0.3°C while the mutant cysteine produces a peak at 84.7±0.3°C. Homozygotes for either allele produce one peak while heterozygotes produce peaks at both T_m_s. All assay plates included two susceptible ORL1952 strain and two resistant PR strain samples as negative and positive controls, respectively. Most plates also contained artificial heterozygotes created by including ORL1952 and PR homogenates in the same sample. Well positions of individual mosquitoes were maintained in both plates (one plate for each locus) to allow genotyping of an individual for both SNPs. Frequencies for each of the nine genotypes (VVFF, VVFC, VVCC, VIFF, VIFC, VICC, IIFF, IIFC, IICC) were calculated by dividing the specific genotype by the total number tested from each area.

**Table 2.**
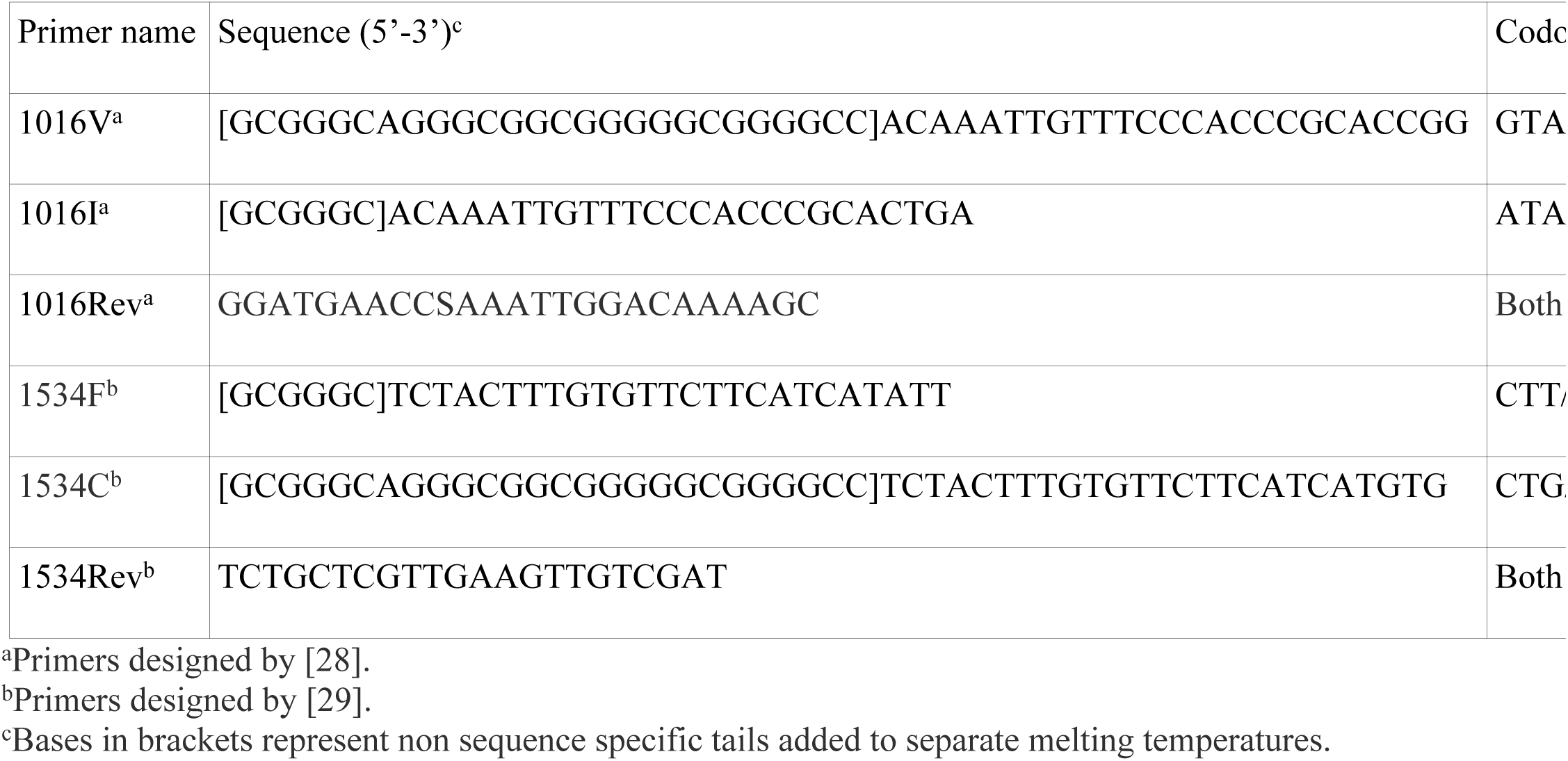
PCR primers used for *kdr* allele analysis.

### Mapping

Base maps in this publication were created using ArcGIS^®^ software by Esri. ArcGIS^®^ is the intellectual property of ESRI and is used herein under license. Copyright © ESRI. GIS data sources were ESRI and Tele Atlas. All rights reserved. (For more information about Esri^®^ software, please visit www.esri.com.’) Permission to publish this content was verified from the ESRI Redistribution Rights Matrix at https://www.esri.com/~/media/Files/Pdfs/legal/pdfs/redistrights106.pdf?la=en

Maps were exported to GIMP2.8 and additional layers with the *kdr* or resistance ratio data from this manuscript was added. Pie chart representations of *kdr* allele frequencies were created with Microsoft excel and exported to layers added to the basemaps.

## Results

### Topical bioassay

All Florida strains of *Ae. aegypti* were resistant to permethrin when compared to the ORL1952 strain, which has been in a continuous laboratory colony for nearly seventy years (Fig 1). The field strains showed varied levels of resistance, from 6-fold to 61-fold compared to the ORL1952 strain. The two most susceptible *Ae. aegypti* strains were collected from flowerpots on Big Coppitt Key (Monroe County) and from oviposition cups in Cortez (Manatee County) at 6.0 and 6.8-fold, respectively. *Ae. aegypti* mosquitoes collected from a tire facility in Orlando (Orange County) were about 17.2-fold more resistant than *Ae. aegypti* collected from the same area generations earlier and used to produce the ORL1952 strain. While most of the FL strains were 15 to 35-fold resistant compared to the lab strains, several strains with higher RR were identified. The strain from New Port Richey in Pasco County had the highest RR at 61.8-fold, which is similar to the RR of the Puerto Rico-resistant reference strain [15]. Strains from Miami Beach and Fort Myers had RRs above 50.

**Figure 1.**
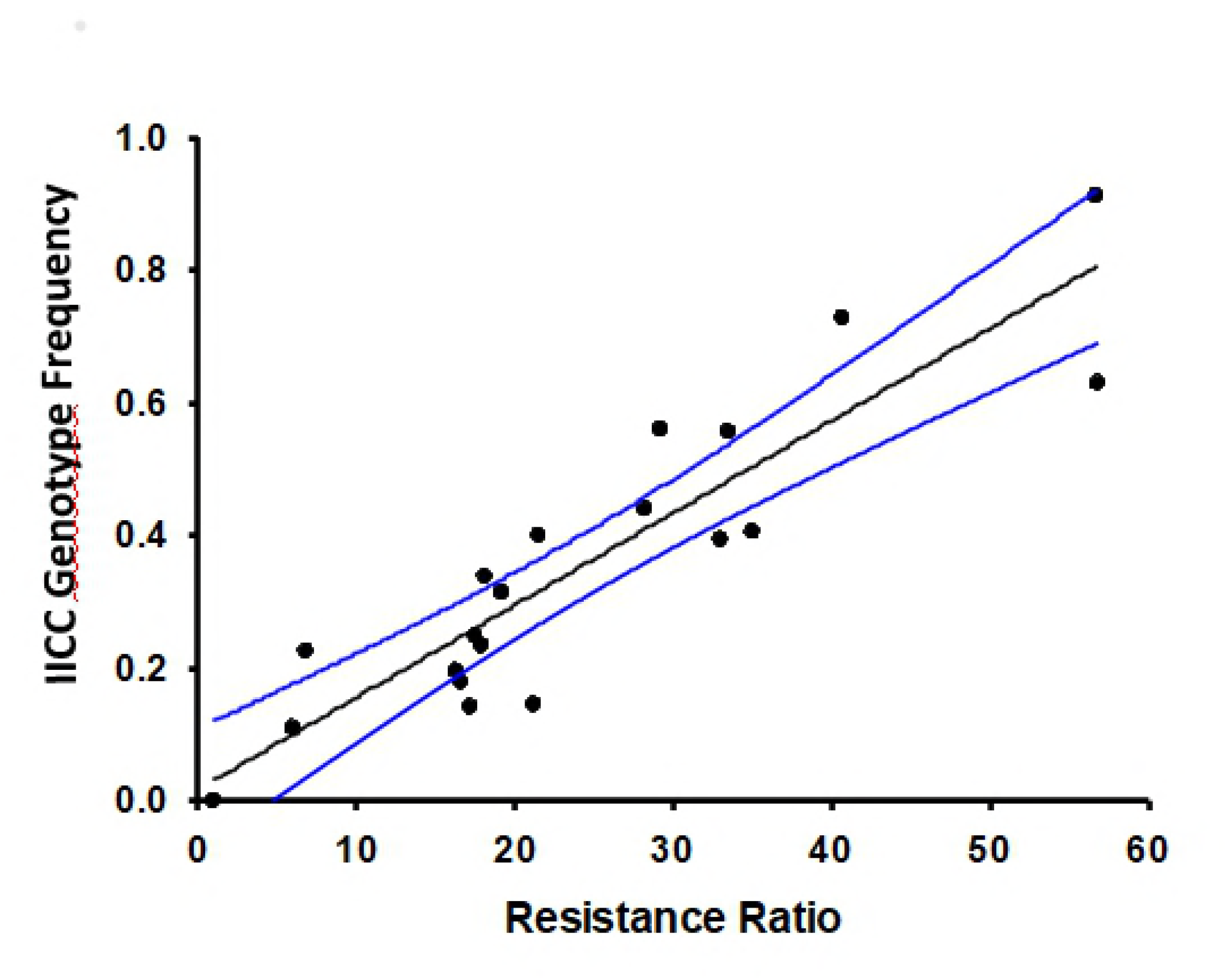
Permethrin resistance ratios of *Ae. aegypti* and *Ae. albopictus* (in blue) strains compared to the susceptible ORL1952 laboratory strain. Base maps were sourced from ESRI and Tele Atlas data through ArcGIS Online under an enterprise license with USDA. Additional layers were added to the base map using GIMP 2.8

The variability we observed in RR throughout the state was also seen at finer resolution within several, but not all, counties. In Miami-Dade County, the Miami Beach and East Wynwood strains were relatively resistant (58-fold and 33-fold), while nearby locations like South Wynwood and central Little River were much more susceptible (Fig 3). Manatee County also had a range of RRs. The Anna Maria Island strain, collected from a densely populated barrier island, had a higher RR than nearby strains from Palmetto or Cortez (Fig 1 and Table 3). We also observed this same trend in Lee County but did not observe universal variation in RR.

**Table 3.** Permethrin resistance ratios of select Florida strains of *Aedes aegypti* and *Ae. albopictus* calculated from direct topical application.

| County | Strain | Species | LD50±SE (ng/mg) | Resistance Ratio |
| --- | --- | --- | --- | --- |
| Lab susceptible | ORL1952 | *Ae. aegypti* | 0.06±0.01 | 1 |
| Manatee | Anna Maria | *Ae. aegypti* | 2.03±0.22 | 35 |
| Manatee | Palmetto | *Ae. aegypti* | 0.77±0.13 | 13.3 |
| Manatee | South Bradenton | *Ae. aegypti* | 1.11±0.18 | 19.2 |
| Manatee | Longboat Key | *Ae. aegypti* | 0.96±0.13 | 16.6 |
| Manatee | Cortez | *Ae. aegypti* | 0.39±0.06 | 6.8 |
| Monroe | Big Coppitt Key | *Ae. aegypti* | 0.35±0.05 | 6 |
| Orange | Orlando | *Ae. aegypti* | 1.34±0.25 | 17.2 |
| Orange | Orlando | *Ae. albopictus* | 0.10±0.02 | 1.3 |
| Seminole | Altamonte Springs | *Ae. aegypti* | 1.42±0.24 | 18.1 |
| Duval | Jacksonville-4 | *Ae. aegypti* | 2.58±0.24 | 33 |
| Hernando | Brooksville | *Ae. albopictus* | 0.12±0.03 | 1.5 |
| Lee | Alva | *Ae. aegypti* | 1.63±0.45 | 16.3 |
| Lee | Alva | *Ae. albopictus* | 0.07±0.05 | 1 |
| Lee | Fort Myers | *Ae. aegypti* | 4.97±0.47 | 56.8 |
| Lee | South Cape Coral | *Ae. aegypti* | 2.90±0.43 | 29.2 |
| Lee | Central Cape Coral | *Ae. aegypti* | 4.05±0.65 | 40.7 |
| Miami-Dade | Wynwood North | *Ae. aegypti* | 2.64±0.57 | 21.5 |
| Miami-Dade | Wynwood Central | *Ae. Aegypti* | 2.15±0.62 | 17.5 |
| Miami-Dade | Wynwood East | *Ae. aegypti* | 4.11±0.64 | 33.5 |
| Miami-Dade | Wynwood South | *Ae. aegypti* | 2.60±0.44 | 21.2 |
| Miami-Dade | Wynwood South | *Ae. albopictus* | 0.17±0.02 | 1.4 |
| Miami-Dade | Little River North | *Ae. aegypti* | 2.19±0.43 | 17.9 |
| Miami-Dade | Little River Central | *Ae. aegypti* | 3.46±1.21 | 28.2 |
| Miami-Dade | Little River Central | *Ae. albopictus* | 0.20±0.02 | 1.6 |
| Miami-Dade | Miami Beach | *Ae. aegypti* | 6.96±1.36 | 56.7 |

*Aedes albopictus* competes with *Ae. aegypti* for oviposition sites, and in several locations *Ae. albopictus* eggs were collected in conjunction with *Ae. aegypti* eggs. *Ae. albopictus* strains reared were also subjected to topical application along with the *Ae. aegypti* counterparts from the same locations (Fig 1, in blue). In the *Ae. albopictus* strains we tested, only slight resistance to permethrin (<2-fold) was observed when compared to the ORL1952 strain. In Miami-Dade County, we observed large differences in RR between *Ae. albopictus* and *Ae. aegypti* even when collected from the same sites in areas such as south Wynwood and central Little River. Duval (Jacksonville), Orange (Orlando), and Lee (Alva) counties also had resistant *Ae. aegypti* and low RR *Ae. albopictus*.

### *Kdr* allele distribution

Examination of *kdr* alleles in *Ae. aegypti* strains from 62 locations showed a range of genotypes (Fig 2 and Fig 3). We observed that most strains were fixed or nearly fixed for the F1534C SNP. For many strains, more than 95% of the tested mosquitoes were homozygous for the 1534C (1534CC) and the remainder of the strain was made up of a few 1534 heterozygotes (1534FC). Only strains from Big Coppitt Key (13% FC), Cortez in Manatee County (38% FC), Central Wynwood (22% FC) and 2 strains from Little River (20% and 14% FC) had more than 10% of mosquitoes that were heterozygous at position 1534. Most strains had no mosquitoes without at least one copy of the 1534C allele but we did find two strains from central Wynwood (8%FF) and Cortez (11% FF) that still had appreciable numbers of susceptible alleles.

**Figure 2.**
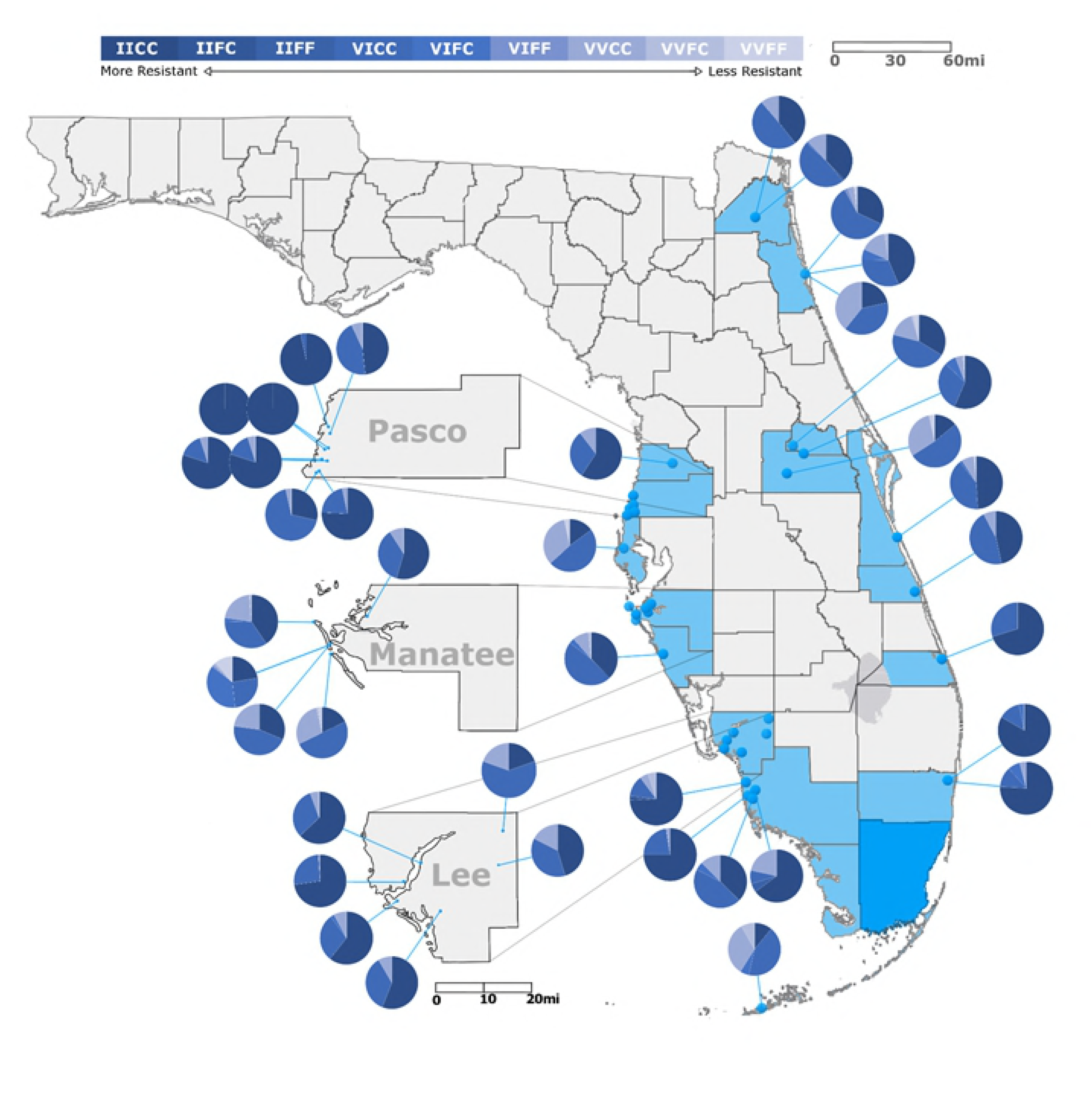
Frequency of *kdr* genotypes in 40 strains of FL *Aedes aegypti*. Genotype frequencies were determined using the methods of [28, 29] as described in the methods section. Specific collection locations that are included in each tested population are noted in Supplemental File 1. Strains included a minimum of 25 individual organisms. Base maps were sourced from ESRI and Tele Atlas data through ArcGIS Online under an enterprise license with USDA. Additional layers were added to the base map using GIMP 2.8. Graphical representation of *kdr* frequencies was produced using Microsoft Excel and included as an additional layer.

**Figure 3.**
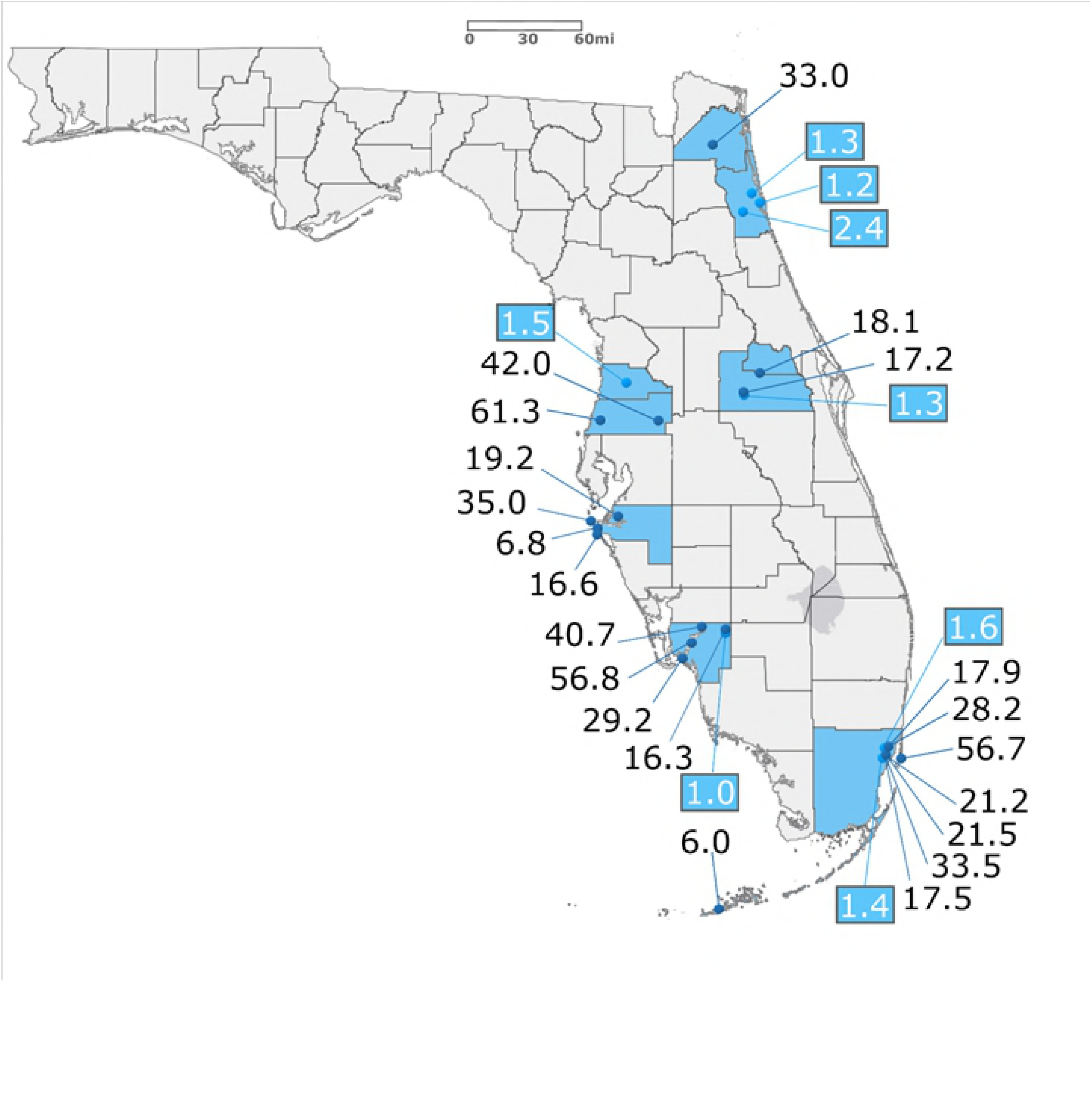
*Kdr* genotype frequencies in 20 populations of *Aedes aegypti* from Miami-Dade County, FL. Genotype frequencies were determined using the methods of [28, 29] as described in the methods section. Specific collection locations that are included in each tested population are noted in Supplemental File 1. All data is based on a minimum of 76 tested individuals. Base maps were sourced from ESRI and Tele Atlas data through ArcGIS Online under an enterprise license with USDA. Additional layers were added to the base map using GIMP 2.8. Graphical representation of *kdr* frequencies was produced using Microsoft Excel and included as an additional layer.

There was much more variation throughout the state at position 1016. Strains from Big Coppitt Key, Longboat Key, Orlando and Clearwater had the lowest percentages of the homozygous 1016II at 11.0, 14.1, 14.9, and 17.8%, respectively. We did observe strains with high levels of 1016II. Six of eight strains examined from Pasco County had 1016II frequencies above 75%, including two strains from New Port Richey at 100% 1016II. Due to the 2016 Zika outbreak and resulting massive public health response, we heavily sampled strains from southeast Florida for allele frequencies. Two strains from Broward and several from Miami-Dade County had 1016II frequencies above 75% including a Miami Beach strain at 91% 1016II (Fig 3). Select strains from Lee and Collier counties in southwest Florida also were above 75% 1016II (Fig 2).

As with RR, in several counties we observed large differences in allele frequency from one part of the county to another. In Lee County (Fig 2), an inland strain from Alva had a relatively low level 1016II frequency (20%) while four strains from near the Gulf Coast all had 1016II frequencies of greater than 55%. This variation was also observed at much closer scale between strains in neighboring cities. The Manatee County Anna Maria Island strain had a much higher 1016II allele frequency than nearby strains from Palmetto or Longboat Key (Fig 2). While overall IICC frequencies were high in Pasco County, we did find variation at the neighborhood level in south Pasco County, where a strain with 75% 1016II was found just blocks from a strain with 30% 1016II (Fig 2). In Miami-Dade, we were able to test four strains from Wynwood and three from Little River collected during the 2016 Zika response (Fig 3). Again, we observed variations in the levels of 1016II within each neighborhood. North Wynwood was slightly more than 25% IICC while south Wynwood was nearly 50%. In Little River, the four strains ranged from 20 to 60%. Just across the water from Wynwood, the Miami Beach strain was greater than 90% IICC.

### Correlating *kdr* genotype with resistance ratio

We regularly observed only six genotypes in the field (Table 4). Only a combined seven mosquitoes of the 4,810 analyzed in this study had genotypes of IIFF, IIFC, or VIFF, which would require an allele coding for a resistant isoleucine at 1016 (1016I) and a susceptible phenylalanine at 1534 (1534F) to be contributed from at least one parent. We did commonly observe the reverse where homozygous or heterozygous susceptible 1016V alleles paired with the homozygous resistant 1534C allele (VVCC or VICC). Plotting the combined dataset for strains with both *kdr* genotype frequencies and resistance ratios from topical application of permethrin showed a strong correlation between increasing RR and increasing 1016II frequency (Fig 4, Pearson correlation coefficient=0.905, P<0.00001, n=20). Strains, like Miami Beach, Fort Myers, and New Port Richey, with high frequencies of 1016II, which, based on the rarity of IIFF and IIFC mosquitoes, implies the genotype IICC were the strains with the highest resistance ratios. In contrast, the strain from Big Coppitt Key had the lowest 1016II frequency and the lowest RR even though the 1534CC frequency was relatively high.

**Table 4.**
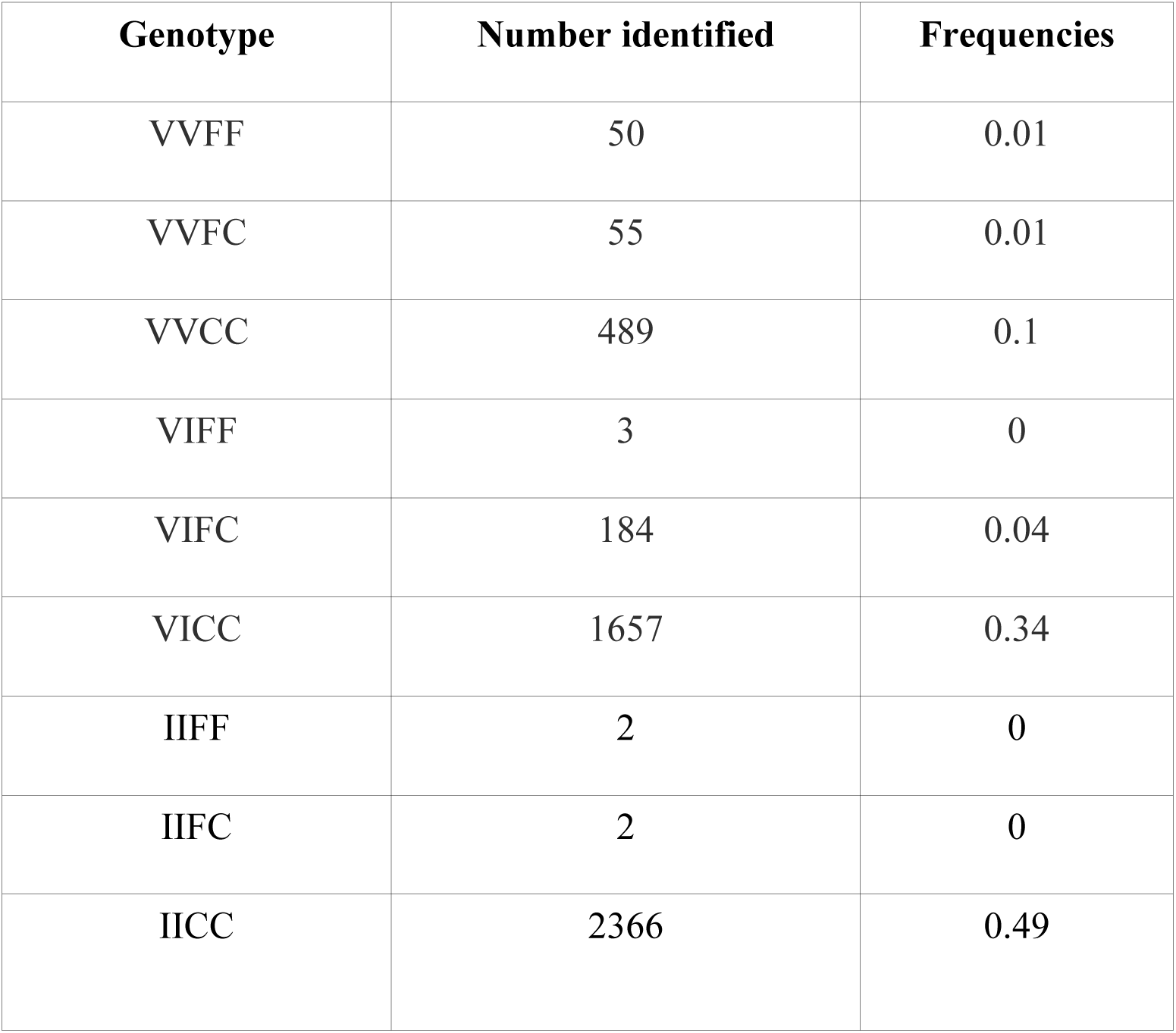
Combined genotype frequencies for 1016 and 1534 alleles in 4,810 Florida *Ae. aegypti*.

**Figure 4.**
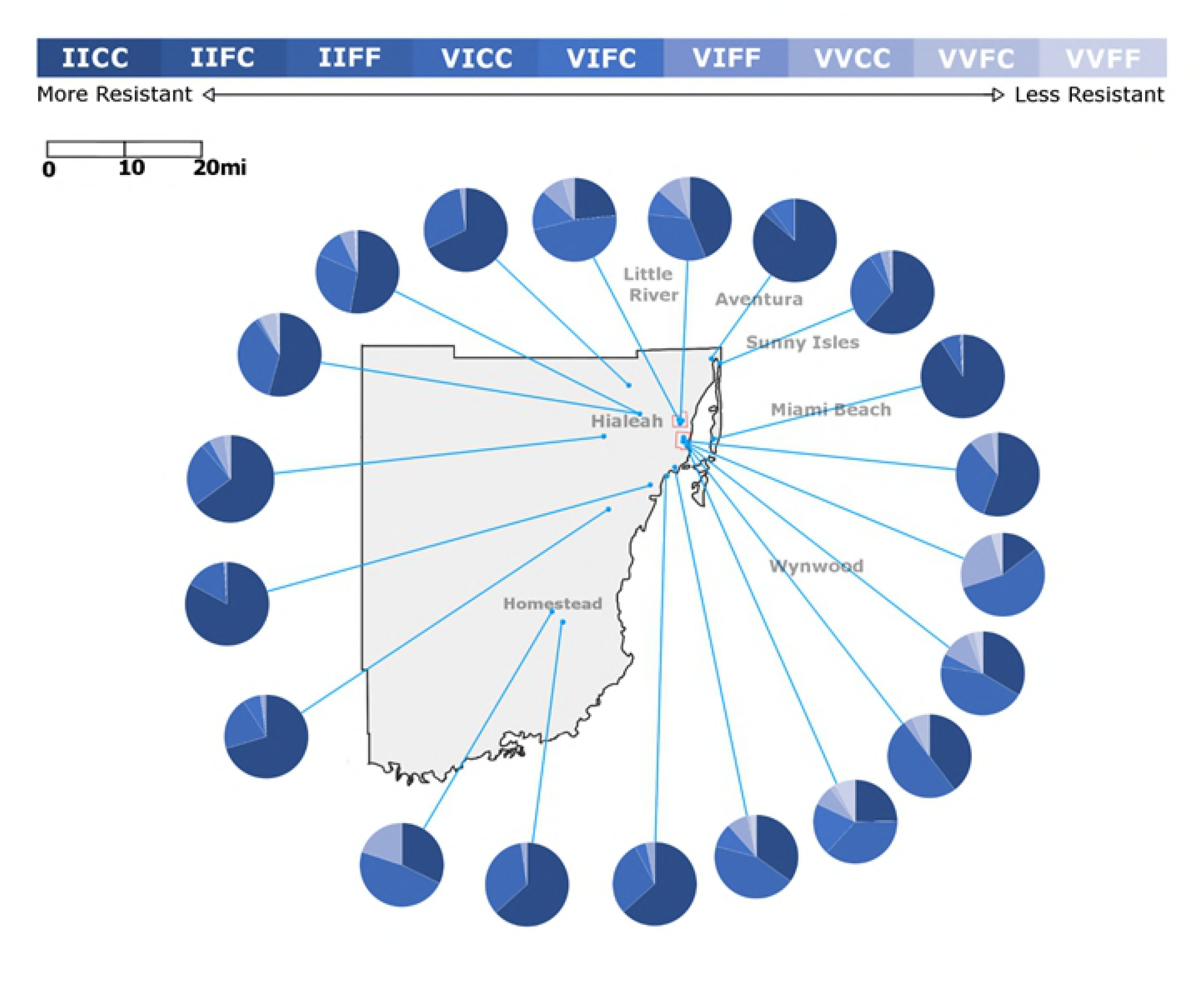
Regression of genotype IICC frequency to resistance ratio. Plot of RR versus IICC frequency indicates a strong correlation between the two factors (Pearson’s Correlation P=0.905). Comparison of RR to other genotypes did not indicate a strong correlation (not shown).

## Discussion

Recent local transmission of Zika virus during 2016 and small outbreaks of dengue virus in 2010 and 2011 demonstrate that an effective IVM plan that attacks multiple life stages to reduce mosquito numbers is a necessity. However, when disease transmission is active, chemical adulticiding can be the only means to immediately reduce the population of potentially infected mosquitoes. While there is some debate as to overall efficacy of adulticiding against *Ae. aegypti* and *Ae. albopictus*, there is little question that it is a necessary part of the response that allows other slower methods of control like larviciding and source reduction to gain a foothold. Pyrethroids are the major class of chemical adulticides that Florida mosquito control programs use operationally. Thus, the efficacy of pyrethroids on *Aedes* vectors is a critically important part of a control program.

Direct topical application of permethrin clearly demonstrated that resistance was widespread in *Ae. aegypti* strains throughout Florida. RRs ranged from about 6-fold in strains from Manatee County and Big Coppitt Key to approximately 60-fold in Miami Beach, Cape Coral, and New Port Richey. Genotyping thousands of individuals indicated that common *kdr* alleles were also widely distributed in the state. While these findings are noteworthy as they represent the first published report of widespread pyrethroid resistance and *kdr* alleles in the southeast US, this result is not surprising and could be predicted based on the results from other locations including Puerto Rico, Mexico and, more recently, in invasive CA *Ae. aegypti* strains [9, 11, 18].

The dataset described in this study does reveal wide variations in both RR and *kdr* alleles within small geographic areas. Examining Miami-Dade County, the strain collected from Miami Beach was highly resistant and had high levels of *kdr*. However, strains from inland Wynwood and Little River were much less resistant than the Miami Beach strain. Even strains from the opposite ends of neighborhoods differed. We saw this disparate pattern in south Miami-Dade, Manatee, and Lee counties. In Pasco County, we saw very different genotypes in strains geographically close to one another, separated only by a major highway. Considering this variability, along with the relatively short flight ranges and limited immigration in *Ae. aegypti* [30], the very different allele frequencies we observed in geographically close strains would support performing testing in numerous areas of a control district to get an accurate resistance picture.

In contrast to the resistance in the *Ae. aegypti* strains, we observed very little permethrin resistance in *Ae. albopictus* statewide. This was true whether *Ae. aegypti* were present or absent from the same collections. Waits et al. [22] showed very low levels of resistance in St. Johns County strains collected from areas without *Ae. aegypti*. Miami-Dade, Orange, and Lee counties all had the same pattern of resistant *Ae. aegypti* and much less resistant *Ae. albopictus*. It has been proposed that the development of pyrethroid resistance is much more difficult in *Ae. albopictus* due to sequence differences in the voltage-gated sodium channel (NaV) that make *kdr* much less likely, although recent reports indicate it may be possible [31, 32]. Our laboratory efforts to induce permethrin resistance in wildtype Florida *Ae. albopictus* by gentle pressuring have failed. At this time, pyrethroid resistance in Florida *Ae. albopictus* does not appear to be an issue that could lead to adulticide failure.

An important observation made due to this work is the correlation between increasing RR and the frequency of the IICC genotype in *Ae. aegypti*, which has been anecdotally observed in other studies using the CDC bottle bioassay [6, 9, 11, 33]. The linkage between *kdr* mutations and a strong correlation with resistance ratios has also been observed in other dipterans [34, 35]. While more research must be done to validate the correlation, this dataset adds another 20 strains with both resistance and *kdr* data that support using *kdr* genotype as a surrogate to estimate pyrethroid resistance levels in mosquitoes [36]. The use of allele frequencies has several potential benefits. Genotype data are relatively easy to collect, the results are produced in hours, and dead mosquitoes collected during standard surveillance activities can be used to provide information on resistance. With limited budgets and personnel at the operational vector control district level, predictive estimation of resistance levels could produce useful operational data from activities already being done without requiring additional efforts to collect or produce mosquitoes for bioassay testing.

Nearly a decade after Donnelly et al. [36] asked whether *kdr* genotypes are predictive in mosquitoes there is, to our knowledge, no published study that shows a pyrethroid-resistant strain of *Ae. aegypti* without also showing the presence of *kdr* alleles. We suggest that the major benefits to be gained from use of allele frequencies as an estimator of resistance would be improvements in coverage area (this study, 62 strains with allele frequencies vs. 26 by direct topical application) and better operational decision-making as vector control programs could access more timely information on area specific resistance levels. These challenges of getting wide coverage and providing this information on an operationally useful timeline have been present in recent response efforts to Zika virus. The efforts of CDC and vector control units in Puerto Rico and the efforts of the authors and others in Florida to develop wide-area pyrethroid resistance maps by relying strictly on bioassay data show that it is currently a slow, labor intensive process. In Florida at least, the epidemic had passed by the time more than a few strains had been tested by bioassay. Until there is reliable, published evidence to argue against the use of *kdr* frequencies as a predictor, it is at present the only rapid way to assess strains across a large area.

This study shows that permethrin resistance is widely present in variable intensity in *Ae. aegypti* throughout Florida. This variability points to the need to include resistance testing as part of an IVM plan as well as examine resistance in more than one location. But how do we use this resistance information to improve vector control? Clearly, the strain of *Ae. aegypti* in Miami Beach is very different from nearby Wynwood and would likely call for different treatment strategies. A treatment with permethrin would likely have much less effect in Miami Beach in comparison to other less resistant locations.

Our study also points to the value of a collaborative approach from motivated stakeholders to develop resistance information. The state of Florida began regular conferences to bring together vector control districts, researchers, and public health resources months before the first locally transmitted case of Zika was reported in 2016. Like the CDC response in Puerto Rico, development of resistance information was an early and ongoing part of this process. The dataset in this study represents the result of thousands of hours of effort from vector control districts, vector control contractors, the state of Florida, and the federal government to produce operationally useful resistance information to protect public health and improve the efficacy of control operations. However, even with these efforts, this study is very limited in scope. We examined only permethrin resistance and although the literature shows the patterns we observed would likely be applicable to other pyrethroids [11, 15], work to define statewide patterns of resistance to synergized products or organophosphates still needs to be addressed.

## Supporting Information Legends

**Supporting File 1. *Collection information and kdr data for strains used in this study***. This excel file contains detailed location information on collection sites, lifestages tested, numbers of organisms tested and raw *kdr* allele data.

